# Capacity of countries to reduce biological invasions

**DOI:** 10.1101/2021.02.04.429788

**Authors:** Guillaume Latombe, Hanno Seebens, Bernd Lenzner, Franck Courchamp, Stefan Dullinger, Marina Golivets, Ingolf Kühn, Brian Leung, Núria Roura-Pascual, Emma Cebrian, Wayne Dawson, Christophe Diagne, Jonathan M. Jeschke, Cristian Perez-Granados, Chunlong Liu, Dietmar Moser, Anna Turbelin, Piero Visconti, Franz Essl

## Abstract

The extent and impacts of biological invasions on biodiversity are largely shaped by an array of socio-ecological predictors, which exhibit high variation among countries. Yet a global synthetic perspective of how these factors vary across countries is currently lacking. Here, we investigate how a set of five socio-ecological predictors (Governance, Trade, Environmental Performance, Lifestyle and Education, Innovation) explain i) country-level established alien species (EAS) richness of eight taxonomic groups, and ii) country capacity to prevent and manage biological invasions and their impacts. Trade and Governance together best predicted the average EAS richness, increasing variance explained by up to 54% compared to models based on climatic and spatial variables only. Country-level EAS richness increased strongly with Trade, whereas high level of Governance resulted in lower EAS richness. Historical (1996) levels of Governance and Trade better explained response variables than current (2015) levels. Thus, our results reveal a historical legacy of these two predictors with profound implications for the future of biological invasions. We therefore used Governance and Trade to define a two-dimensional socio-economic space in which the position of a country captures its capacity to address issues of biological invasions. Our results provide novel insights into the complex relationship between socio-ecological predictors and biological invasions. Further, we highlight the need for designing better policies and management measures for alien species, and for integrating biological invasions in global environmental scenarios.

## Introduction

The proliferation of alien species – i.e. species that are intentionally or unintentionally introduced by humans in regions beyond their native ranges – has become a signature of human-induced global environmental change. A substantial proportion of these species becomes a permanent addition to regional biota (established alien species – EAS hereafter), and a subset of these species, known as invasive alien species (IAS), are a leading cause of biodiversity decline (1–3) and can adversely affect human livelihoods (4–7). Globally, the number of EAS has been steadily increasing in recent decades, and this trend does not show any sign of saturation (8). Meanwhile, the current state and particularly the future trajectories of EAS impacts remain highly uncertain (9, 10). Still, there is a distinct lack of consideration for the impacts of biological invasions in developing long-term global biodiversity conservation frameworks and scenarios (11, 12).

Environmental and economic factors have been repeatedly demonstrated to be important predictors for biological invasions at the global scale (13–16). Additionally, experts also consider political, social and technological factors to be important (10, 17). For example, countries with low Human Development Index (18) are severely constrained in their capacity to manage biological invasions and mitigate their impacts (19). In addition, the relationship between governance and biological invasions is complex, as countries with high levels of governance (i.e. countries in which governments are selected, monitored and replaced democratically, in which governments can effectively formulate and implement sound policies, and in which citizens and the state respect the institutions that govern economic and social interactions among them (20)) are more susceptible to biological invasions if they also have high per capita-GDP (21). Low levels of governance and high levels of corruption have also been associated with higher exports of alien species, as regulations of outbound pathways are poorly implemented and subsequently lead to greater potential rates of introduction in importing countries (22). In many instances, species are also deliberately released because of their perceived or realized economic benefits. For instance, plants of the genera *Melinis* and *Urochloa* were released in Brazil as livestock feed, but are now known to fundamentally alter the ecosystems in which it is established via dominance and changes in fire regimes (23). In response to growing threats from biological invasions, many countries with high richness of alien species have expanded and implemented new legislations on alien species since the 1990s (24). Quantitative analyses are nonetheless scarce for political, social and technological predictors.

Understanding how socio-ecological predictors together shape the current and future state of biological invasions at the country scale is crucial to design and implement efficient policies and future global scenarios for biological invasions (11, 17). Recent global studies considering the combined role of social, political, environmental and socio-economic predictors for the future of biological invasions have mostly relied on expert knowledge (10, 25). Therefore, there is a need for a comprehensive quantitative assessment of these relationships.

Here, we compare 125 countries (excluding some regions separate from mainland, which can have different invasion dynamics) against a set of political, economic, environmental, social and technological predictors, which are considered to be essential drivers of biological invasions (10). For each country, we quantify current and – if available – historical conditions using five predictors (Governance, Trade, Environmental performance, Lifestyle and Education, and Innovation; Table 1). We i) examine the relationships between these predictors and then relate their ii) current (2015) and iii) past (1996) values to EAS richness per country. As a response variable, we use country-level EAS richness of eight taxonomic groups (plants, ants, amphibians, reptiles, fishes, birds and spiders) based on the most comprehensive country-level data set on EAS richness (15). Moreover, we relate these predictors to the national response capacities to manage and mitigate biological invasions and their impacts presented in (19).

**Table 1.** Main predictors of biological invasions (as identified by (17)) and their corresponding descriptor variables. Note that variables with * were discarded from analyses (see Methods).

| Predictor | Relationship with biological invasions | Variable | Description | Source |
| --- | --- | --- | --- | --- |
| Governance | The political context influences the capacity of a country to vote and apply appropriate policies and management actions to control and prevent the introduction of IAS. | Rule of Law | Perception of the extent to which the population has confidence in and abides by the rules of society, and in particular the quality of contract enforcement, property rights, the police, and the courts, as well as the likelihood of crime and violence. | The World Bank (36) |
|  |  | Government Effectiveness | Perception of the quality of public services, the quality of the civil service and the degree of its independence from political pressures, the quality of policy formulation and implementation, and the credibility of the government's commitment to such policies. | The World Bank (36) |
|  |  | Voice and Accountability | Perception of the extent to which a country's citizens are able to participate in selecting their government, as well as freedom of expression, freedom of association and a free media. | The World Bank (36) |
|  |  | Control of Corruption | Perception of the extent to which public power is exercised for private gain, including both petty and grand forms of corruption, as well as "capture" of the state by elites and private interests. | The World Bank (36) |
|  |  | Political Globalization* | Summary of the diffusion of countries' government policies and the ability to engage in international political cooperation. | KOF Swiss Economic Institute (37) |
| Trade | The importation of goods and services into a country facilitates the | Imports in Good and Services | Value of all goods and other market services received from the rest of the world. | The World Bank (36) |
|  |  | Per capita Gross National | Sum of a country's gross domestic product (GDP) plus | The World Bank (36) |

|  | introduction of propagules. | Income (GNI)* | net income (positive or negative) from abroad, divided by population size. |  |
| --- | --- | --- | --- | --- |
| Environment | Environmental conditions, including disturbance, land cover, pollution, etc., can influence the establishment of alien species. | Environmental Performance Index (EPI) | Summary of the state of sustainability of countries, based on 32 performance indicators across 11 issue categories (Biodiversity and habitat, Ecosystem services, Fisheries, Water resources, Climate change, Pollution emissions, Agriculture, Waste management, Heavy metals, Sanitation and drinking water and Air quality). | (50) |
| Lifestyle and Education | People's lifestyle, their connections with other cultures and therefore geographical areas, itself influenced by education, can move and introduce propagules to novel environments. Lifestyle and education are also likely linked to other predictors, such as governance. | Average level of education of the population<br>Information Globalization Index (de jure)<br>Cultural Globalization Index (de jure) | Average of the maximum level of education across countries' inhabitants, using a scale from 1 to 5 to quantify levels of education.<br>Summary of countries' ability to share information with other countries.<br>Summary of countries' openness towards and the ability to understand and adopt foreign cultural influences | European Commission & Joint Research Centre (40)<br>KOF Swiss Economic Institute (37)<br>KOF Swiss Economic Institute (37) |
| Innovation | Technological innovations can offer means to control and prevent the introduction of alien species, but may also facilitate trade activities and contribute to impact the environment. | Global Innovation Index | Summary of countries' capacity for, and success in, innovation, based on variables from multiple sources. | Cornell University, INSEAD, WIPO (39) |

Based on the results from these analyses, we show how Governance and Trade can be used to identify a two-dimensional, socio-economic space describing the capacity of countries to mitigate alien species spread and impact. We assess how different geopolitical groups of countries (Figure S1) perform in this socio-economic space. Finally, we show how countries and geopolitical regions have changed their position in this socio-economic space since 1996, and explain why divergences between country trajectories are crucial to capture the main challenges they are currently facing to tackle invasions.

## Results

### Relationships between predictors

Governance (quantified as the mean of Rule of Law, Government Effectiveness, Voice and Accountability and Control of Corruption; Table 1), Environment (measured by the Environmental Performance Index, which includes land use; Table 1) and Lifestyle and Education (quantified as the mean of the average level of education of a population, the Information Globalization Index and the Cultural Globalization Index; Table 1) were highly correlated (0.77 ≤ r ≤ 0.85) across countries, and less so with Trade (measured as total imports in Good and Services; r ≤ 0.59). Innovation (measured by the Global Innovation Index) was moderately correlated with all other predictors (0.59 ≤ r ≤ 0.62). A principal component analysis (PCA) confirmed the distinction between these three groups of predictors (Figure S2g).

### Predictors and numbers of established alien species in countries

Using recent (2015) data on predictors, all mixed-effects models using only one of the five predictors (in addition to climatic and spatial variables, including mean annual temperature, mean annual precipitation, country area, sampling effort, mainland / inland status and broad geographical region; see Methods) significantly explained observed overall richness of EAS, increasing marginal r^2^ values by between 8% and 25% in absolute values (between 15% and 46% relative increase) compared to models only including climatic and spatial variables (Table 2). Model comparison using the Akaike Information Criterion corrected for small-sample size (AICc) revealed that Trade was the best predictor of richness for overall EAS data and for most individual taxonomic groups, i.e. plants (together with Governance), ants, amphibians, reptiles, fishes and birds (Table 2, Figure S3). Meanwhile, for mammals and spiders, Lifestyle and Education was the most important predictor. The effect size varied between taxonomic groups, being mostly null for ants and representing almost half of the marginal variance for fishes. The relationships between several predictors and EAS richness were non-linear for most taxa. For Trade, the relationship was positive quadratic for most taxonomic groups, indicating an acceleration of EAS richness as Trade increased (the relationship was cubic, slightly decelerating at high Trade values for birds). For Innovation, the relationship was also quadratic and accelerating for all taxa. In contrast, the relationships between EAS richness and Governance, Environmental Performance and Lifestyle and Education were either quadratic or cubic and tended to decrease at high values (Table 2, Figure S3).

**Table 2.** Results of model fitting for explaining EAS richness and national capacities in 125 countries based on the small-sample size corrected Akaike Information Criterion (AICc) for 2015. In bold are the models with the lowest AICc values. The ΔAICc is the difference with the lowest AICc values over all predictors and all polynomial forms. r^2^ values are the marginal variance. Values in the first column are those when only environmental predictors and random effects were included, and all predictors were excluded (the marginal r^2^ is therefore 0 for the national capacity models). Values between brackets indicate the gains in marginal r^2^ compared to models with no predictor.

| Response variable | Governance | Trade | Environment | Lifestyle and Education | Technology |
| --- | --- | --- | --- | --- | --- |
| All taxa combined<br>$r^2 = 0.54$<br>ΔAICc = 48.1 | Quadratic<br>$r^2 = 0.68$<br>(+0.14)<br>ΔAICc = 44.06 | **Quadratic**<br>$r^2 = \mathbf{0.79}$<br>**(+0.25)**<br>ΔAICc = **0.00** | Quadratic<br>$r^2 = 0.66$<br>(+0.13)<br>ΔAICc = 42.55 | Quadratic<br>$r^2 = 0.64$<br>(+0.10)<br>ΔAICc = 46.28 | Quadratic<br>$r^2 = 0.62$<br>(+0.08)<br>ΔAICc = 47.22 |
| Plants<br>$r^2 = 0.45$<br>ΔAICc = 8.94 | Cubic<br>$r^2 = 0.48$<br>(+0.02)<br>ΔAICc = 6.00 | **Quadratic**<br>$r^2 = \mathbf{0.54}$<br>**(+0.09)**<br>ΔAICc = **0.00** | Cubic<br>$r^2 = 0.52$<br>(+0.06)<br>ΔAICc = 5.97 | Quadratic<br>$r^2 = 0.54$<br>(+0.09)<br>ΔAICc = 2.99 | Quadratic<br>$r^2 = 0.46$<br>(+0.00)<br>ΔAICc = 8.82 |
| Ants<br>$r^2 = 0.66$<br>ΔAICc = 6.58 | Quadratic<br>$r^2 = 0.64$<br>(-0.02)<br>ΔAICc = 5.92 | **Quadratic**<br>$r^2 = \mathbf{0.61}$<br>**(-0.05)**<br>ΔAICc = **0.00** | Quadratic<br>$r^2 = 0.64$<br>(-0.02)<br>ΔAICc = 3.91 | Quadratic<br>$r^2 = 0.68$<br>(+0.03)<br>ΔAICc = 0.80 | Quadratic<br>$r^2 = 0.62$<br>(-0.03)<br>ΔAICc = 4.70 |
| Amphibians<br>$r^2 = 0.47$<br>ΔAICc = 14.97 | Cubic<br>$r^2 = 0.53$<br>(+0.06)<br>ΔAICc = 10.64 | **Quadratic**<br>$r^2 = \mathbf{0.58}$<br>**(+0.12)**<br>ΔAICc = **0.00** | Quadratic<br>$r^2 = 0.54$<br>(+0.07)<br>ΔAICc = 9.60 | Quadratic<br>$r^2 = 0.50$<br>(+0.04)<br>ΔAICc = 12.47 | Quadratic<br>$r^2 = 0.52$<br>(+0.06)<br>ΔAICc = 11.84 |
| Reptiles<br>$r^2 = 0.28$<br>ΔAICc = 26.28 | Cubic<br>$r^2 = 0.35$<br>(+0.06)<br>ΔAICc = 19.58 | **Cubic**<br>$r^2 = \mathbf{0.46}$<br>**(+0.18)**<br>ΔAICc = **0.00** | Quadratic<br>$r^2 = 0.34$<br>(+0.06)<br>ΔAICc = 17.66 | Quadratic<br>$r^2 = 0.28$<br>(+0.00)<br>ΔAICc = 20.94 | Quadratic<br>$r^2 = 0.37$<br>(+0.09)<br>ΔAICc = 16.71 |
| Fishes<br>$r^2 = 0.29$<br>ΔAICc = 43.27 | Cubic<br>$r^2 = 0.40$<br>(+0.11)<br>ΔAICc = 31.15 | **Quadratic**<br>$r^2 = \mathbf{0.56}$<br>**(+0.27)**<br>ΔAICc = **0.00** | Cubic<br>$r^2 = 0.49$<br>(+0.20)<br>ΔAICc = 18.89 | Quadratic<br>$r^2 = 0.43$<br>(+0.13)<br>ΔAICc = 30.18 | Quadratic<br>$r^2 = 0.41$<br>(+0.12)<br>ΔAICc = 28.83 |
| Birds<br>$r^2 = 0.52$<br>ΔAICc = 39.62 | Quadratic<br>$r^2 = 0.61$<br>(+0.08)<br>ΔAICc = 27.34 | **Cubic**<br>$r^2 = \mathbf{0.70}$<br>**(+0.17)**<br>ΔAICc = **0.00** | Quadratic<br>$r^2 = 0.62$<br>(+0.09)<br>ΔAICc = 20.62 | Quadratic<br>$r^2 = 0.62$<br>(+0.10)<br>ΔAICc = 19.44 | Quadratic<br>$r^2 = 0.57$<br>(+0.05)<br>ΔAICc = 30.29 |
| Mammals<br>$r^2 = 0.53$<br>ΔAICc = 12.99 | Quadratic<br>$r^2 = 0.61$<br>(+0.07)<br>ΔAICc = 7.89 | Quadratic<br>$r^2 = 0.60$<br>(+0.07)<br>ΔAICc = 6.56 | Quadratic<br>$r^2 = 0.65$<br>(+0.11)<br>ΔAICc = 0.16 | **Quadratic**<br>$r^2 = \mathbf{0.64}$<br>**(+0.11)**<br>ΔAICc = **0.00** | Quadratic<br>$r^2 = 0.57$<br>(+0.03)<br>ΔAICc = 12.58 |
| Spiders<br>$r^2 = 0.36$<br>ΔAICc = 26.33 | Quadratic<br>$r^2 = 0.47$<br>(+0.11)<br>ΔAICc = 16.23 | **Quadratic**<br>$r^2 = \mathbf{0.59}$<br>**(+0.23)**<br>ΔAICc = **0.00** | Quadratic<br>$r^2 = 0.50$<br>(+0.14)<br>ΔAICc = 11.53 | Quadratic<br>$r^2 = 0.57$<br>(+0.22)<br>ΔAICc = 6.39 | Quadratic<br>$r^2 = 0.44$<br>(+0.08)<br>ΔAICc = 19.94 |
| Proactive national capacity<br>ΔAICc = 28.28 | Quadratic<br>$r^2 = 0.32$<br>ΔAICc = 5.92 | Quadratic<br>$r^2 = 0.16$<br>ΔAICc = 18.89 | Quadratic<br>$r^2 = 0.25$<br>ΔAICc = 17.26 | **Quadratic**<br>$r^2 = \mathbf{0.47}$<br>ΔAICc = **0.00** | Quadratic<br>$r^2 = 0.23$<br>ΔAICc = 15.40 |
| Reactive national capacity<br>ΔAICc = 18.53 | Quadratic<br>$r^2 = 0.11$<br>ΔAICc = 15.26 | **Quadratic**<br>$r^2 = \mathbf{0.26}$<br>ΔAICc = **0.00** | Quadratic<br>$r^2 = 0.16$<br>ΔAICc = 12.62 | Quadratic<br>$r^2 = 0.33$<br>ΔAICc = 2.30 | Quadratic<br>$r^2 = 0.15$<br>ΔAICc = 12.62 |

The combination of Trade and Governance levels of 1996 or averaged over 1996–2015 (i.e. considering both predictors as fixed effects, without interaction) explained EAS richness better than any predictor individually or combining Trade and Governance for 2015 (Figure 1, Table S1). Models only including historical Trade (i.e. 1996 or averaged over 1996–2015) resulted in the best-fitting models for plants and amphibians, and for overall EAS richness. Models using the 2015 data were always worse than models using historical data.

**Figure 1.**
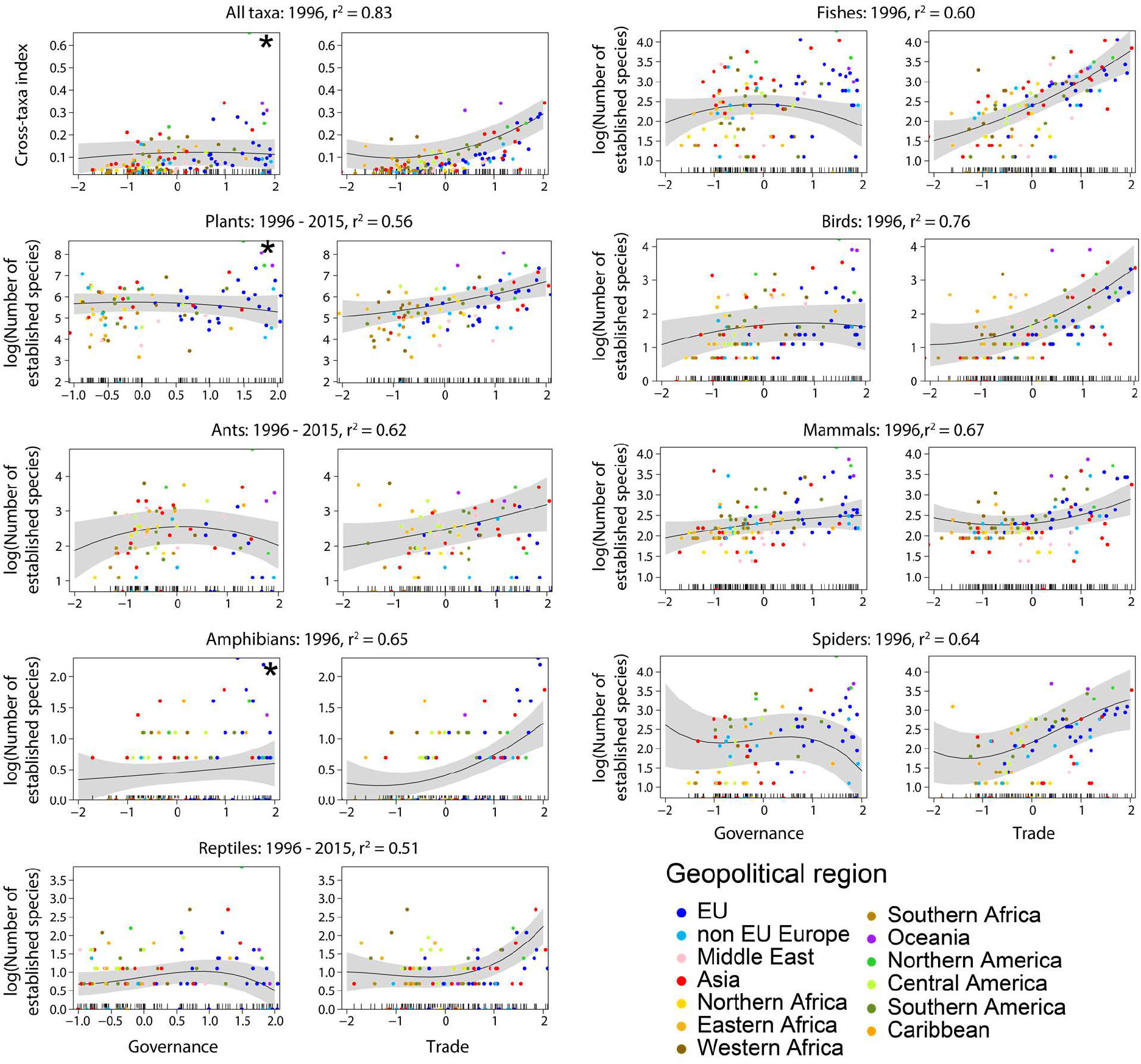
Relationships between the Governance and Trade predictors and the number of EAS in 125 countries, when both predictors were included in linear mixed-effects models. For each taxonomic group, the year or combination of years generating the lowest AICc were selected, and the marginal r^2^ is reported. The number of EAS was controlled for by country area, sampling effort, mean annual temperature and total annual precipitation. Different colors indicate geopolitical regions the countries belong to. Asterisks indicate that a predictor did not improve a model (i.e. a single-predictor model had lower AICc than a two-predictor model).

### Predictors and national response capacities

Lifestyle and Education better explained national proactive capacity to prevent or rapidly respond to emerging IAS than the other predictors for the 2015 data (marginal r^2^ = 0.47; Table 2, Figure S4). The second-best model was the one incorporating Governance (marginal r^2^ = 0.32). For national reactive capacity, i.e. the expertise, resources and willingness to mitigate negative impacts caused by IAS, Trade had the lowest AIC value, but Lifestyle and Education had the highest marginal r^2^ (Table 2). Model performances were higher for proactive than for reactive capacities (Table 2, Figure S4). Quadratic models performed best for all predictors for both types of national capacity, with positive quadratic terms indicating a disproportional strong increase in national capacity with increasing predictor values.

The level of Lifestyle and Education in 2015 also better explained national proactive capacity than any combination of historical predictors (Table S1). Average Governance between 1996 and 2015 was a better predictor than any other model incorporating Governance or Trade. This model showed an acceleration of national proactive capacity with better Governance (Figure 2). Trade for 1996 was the best predictor for reactive capacity and the relationships were linear for most taxonomic groups.

**Figure 2.**
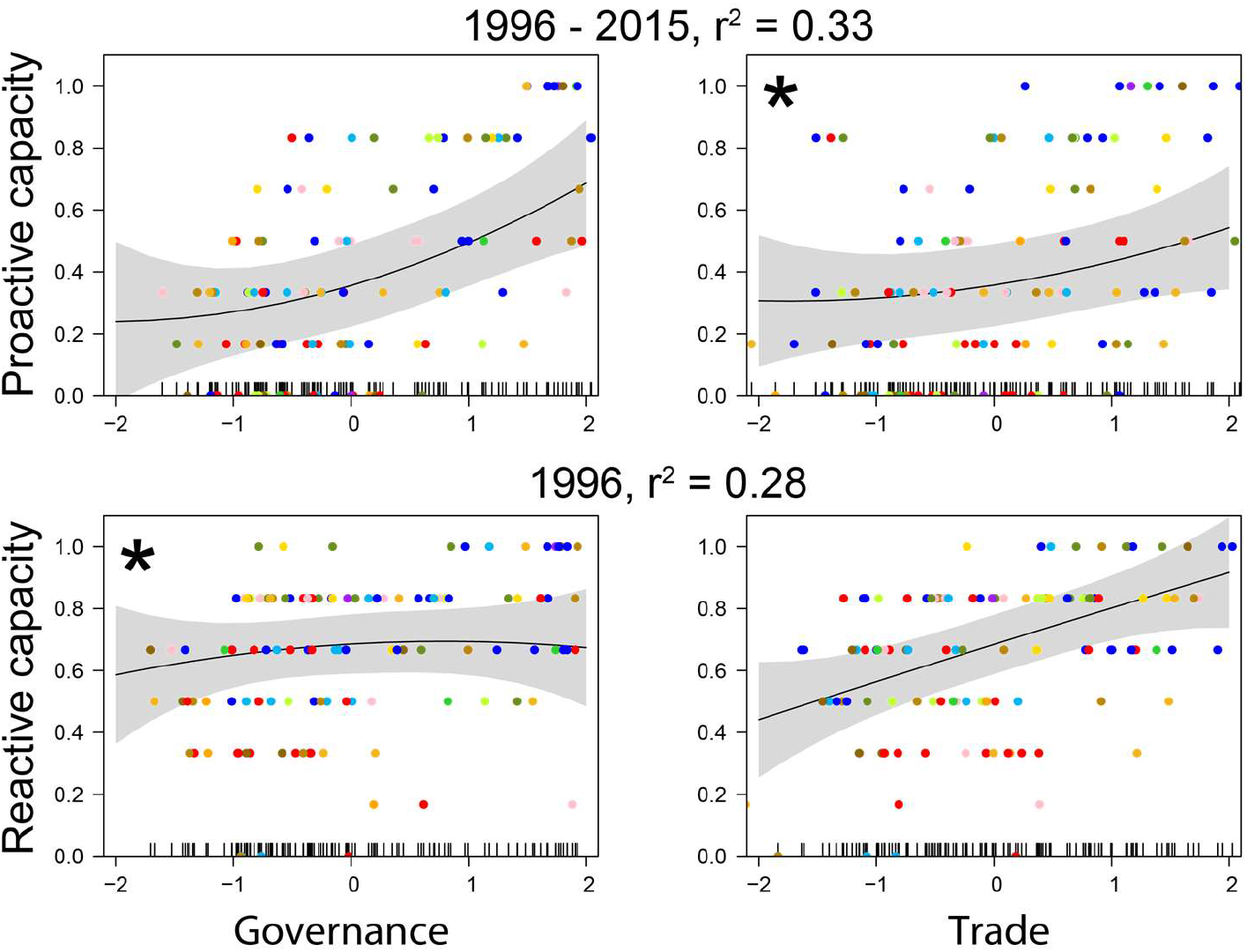
Relationships between Governance and Trade, and national capacities to mitigate the impacts of biological invasions, when both predictors were included in linear mixed models. The year or combination of years generating the lowest AICc were selected, the marginal r^2^ is reported. Different colors indicate the geopolitical regions countries belong to (for legend, see Figure 1). Asterisks indicate that the predictor did not significantly explain established alien species richness (i.e. when linear models with a single predictor generated a lower AIC than when both predictors were included). Rug plots on the inside of the X-axes show the distributions of the data points along individual predictor gradients.

### Mapping countries according to national levels of predictors of invasions

The five predictors selected here were interrelated, but Governance and Trade were the least correlated predictors (Figure S2). Since their historical values were also overall the best (or amongst the best) predictors for both EAS richness and national capacities, we selected Governance and Trade to map countries in a two-dimensional space defined by these two predictors (Figure 3). This two-dimensional approach represents the currently realized socio-economic space of country positions with respect to the main predictors of biological invasion. Contrary to a PCA, whose axes would depend on the data for a given year, using Governance and Trade enables us to assess how countries change their position in time in this fixed socio-economic space (see next section). Capturing country trajectories through time is crucial to understand the dynamics of biological invasions, since they depend on historical legacies.

**Figure 3.**
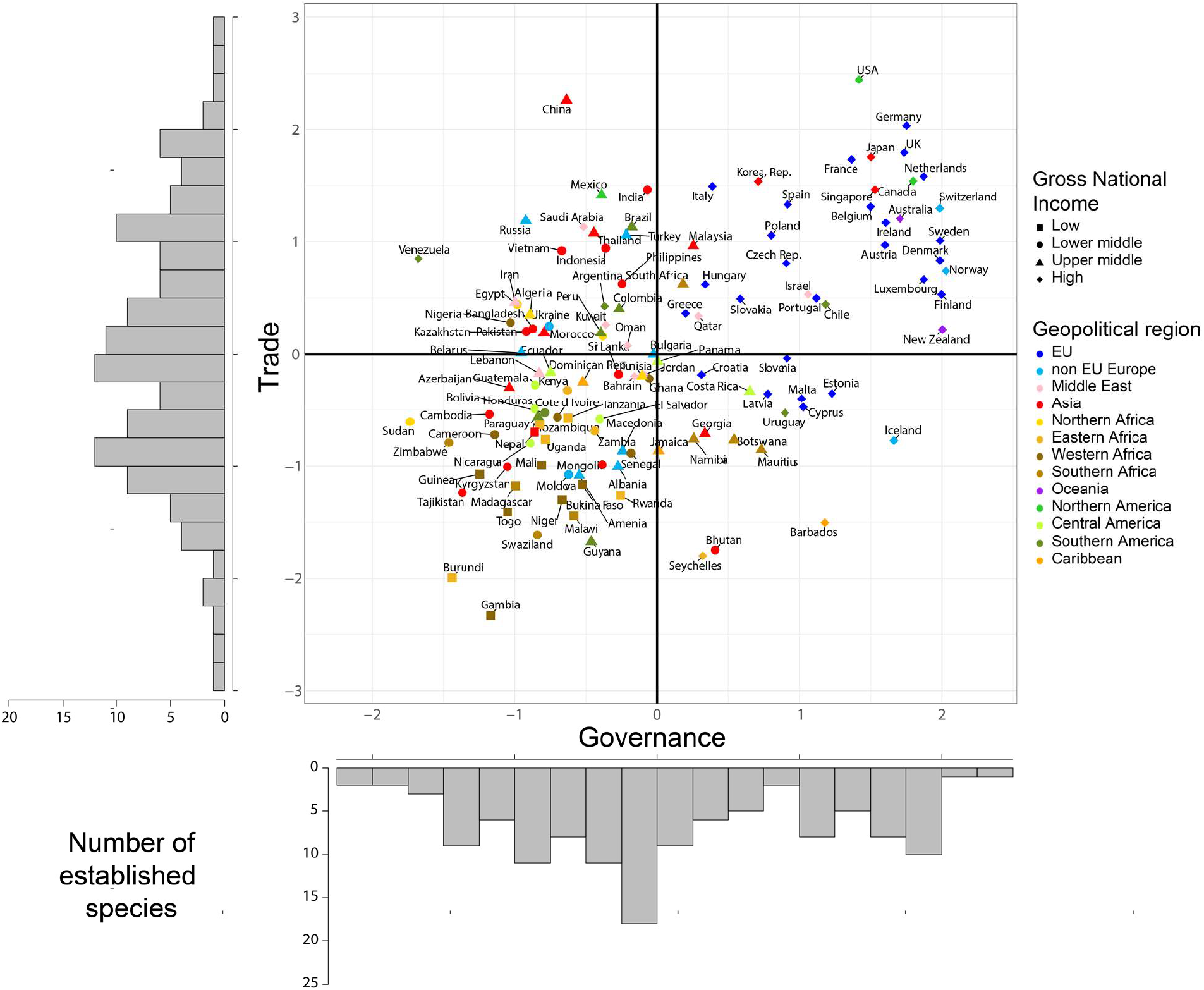
The 125 countries organized in the two-dimensional socio-economic space based on recent (2015) Governance and Trade data. The histograms show the distribution of countries based on Governance and Trade. The bold horizontal and vertical lines indicate the origin axes, which correspond to the centroid of the country distribution. Gross National Income categories are based on the World Bank classification (36).

Consistently with the intermediate, positive correlation between Governance and Trade mentioned above (Figure S2), countries were roughly distributed within an elongated ellipse in the two-dimensional space (Figure 3). Importantly, however, they were not evenly distributed across this ellipse. A cluster analysis revealed that countries can be grouped into four distinct clusters, closely matching the four sectors defined by Governance and Trade (Figure S5). The lower-left section of the socio-economic space contained countries that were characterized by low levels of both Trade and Governance. This section contained the highest number of countries, with 47 out of 125 countries, of which 23 are from the 27 African countries used in our analyses. The upper-right section contained 39 countries with high levels of both Governance and Trade. This category mostly included Western European countries (26 out of 37 countries) and some countries from other continents, including Australia, New Zealand, USA, Canada, Japan and Singapore. The upper-left section contained 23 countries with high levels of Trade but relatively low levels of Governance. This section mostly included Asian countries (9 out of 22 countries). Finally, the lower-right section contained the smallest number of countries (16 countries), which were characterized by low levels of Trade and high levels of Governance. This section contained many island countries. Asian, South-American and African countries were spread over all four sectors, with Asian countries showing the highest variability in their distribution (Figure 3).

### Temporal changes in predictors

Time lag phenomena are common in biological invasions, and our analyses showed that historical data better explained the currently observed EAS richness. We therefore analyzed the trajectories of countries in the two-dimensional socio-economic space defined by Governance and Trade during the past 20 years (Figure 4). All countries have experienced an increase in level of Trade from 1996 to 2018, but changes in Governance were more variable. Countries from continents with high levels of economic development (Australia, Europe and North America) demonstrated high levels of Governance (Figure 4a). Their level of Governance nonetheless tended to increase between 1996 and 2003, and then decreased until 2018. It was even lower in 2018 than in 1996 for Northern America (−0.11 in our standardized scale for this predictor). Governance in Northern African countries has remained at a low level over this period. In contrast, West and East African countries started at a similar level as Northern African countries but saw the second and third largest increase in their level of Governance over time (+0.17 and +016), especially after 2013 for West Africa. Asian countries experienced the largest increase in their level of Governance on average (+0.18). European countries that are not members of the EU experienced a moderate increase (+0.1). Asian countries in the Middle-East saw a rapid increase in the level of Governance between 1996 and 2000 (+0.32), with stable levels of Trade. After 2000, this trend reversed, with a stagnation in the level of Governance and an increase in the level of Trade. Middle Eastern, Caribbean, and especially Southern African countries saw the largest declines in their levels of Governance on average (−0.13, −0.18 and −0.27, respectively).

**Figure 4.**
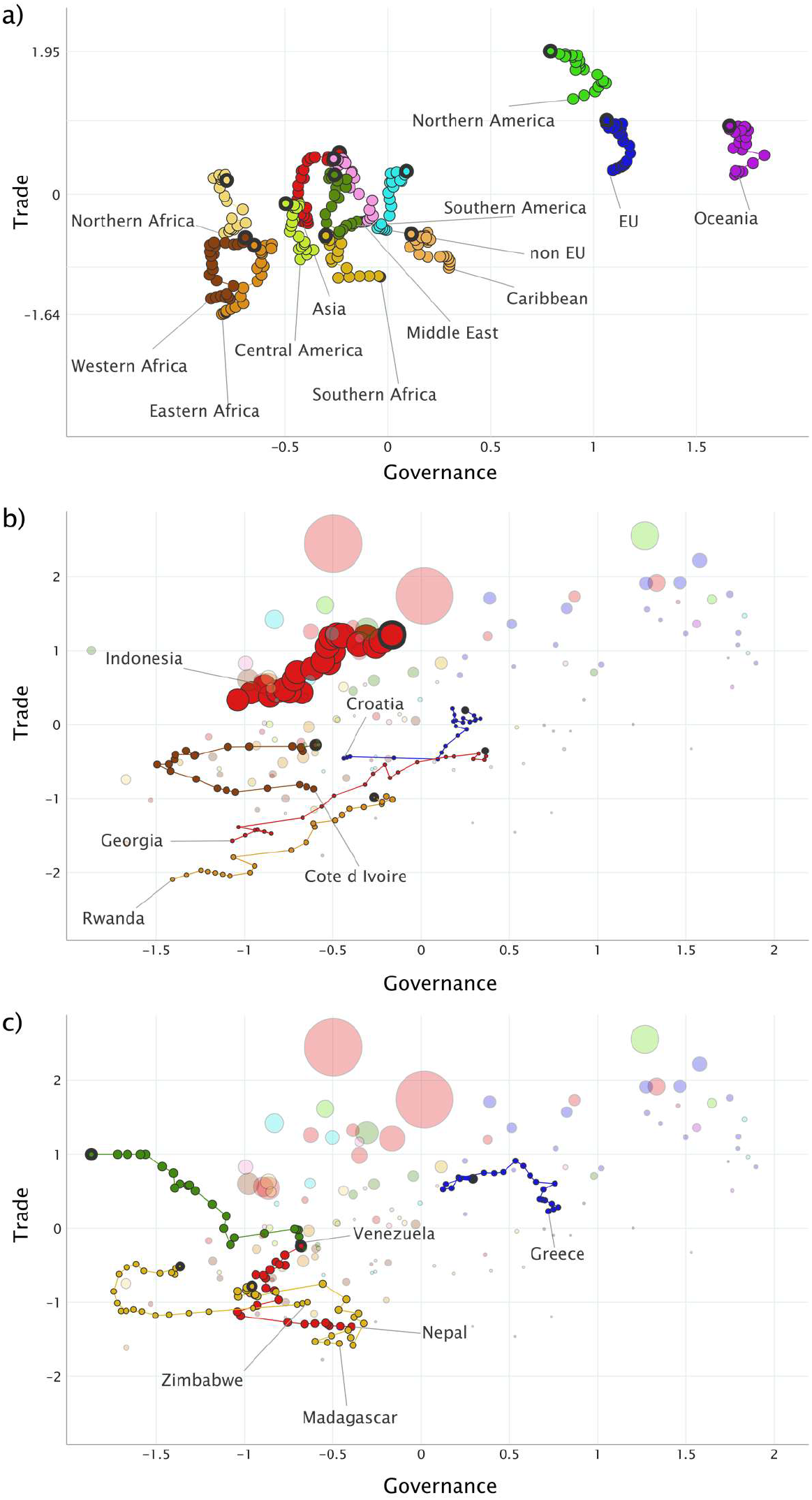
Changes in Governance and Trade between 1996 and 2018 for 125 countries. a) Average changes for main geopolitical regions of the world. b) Changes for countries with the largest increase of Governance between 1996 and 2018. c) Changes for countries with the largest decrease of Governance between 1996 and 2018. Region and country names point towards positions in 1996, and thick bubbles represent positions in 2018. Bubble size illustrates human population size. Visualizations were created using Gapminder (www.gapminder.org).

Results were much more heterogeneous at the country level, with some countries having large increases, decreases or fluctuations in their levels of Governance (Figure 4b,c). Overall, countries with high levels of Governance in 1996 mostly remained close to their initial level. In contrast, countries with intermediate or low levels of Governance changed in either direction. Georgia had the largest increase in level of Governance worldwide (+1.44; Figure 4b), whereas Venezuela had the largest decline (−1.18). Zimbabwe had the second largest decline over only 13 years (i.e. its decline was larger in 13 years than for any other country in 22 years; −1.1), then increased again, but without reaching its former level of Governance (Figure 4c). Ivory Coast had a similar trajectory, but it recovered better and ended with a higher level of Governance in 2018 than in 1996, making the third largest increase in only 13 years (+0.9).

## Discussion

### Socio-ecological predictors of biological invasions

Here, we provide the first comprehensive quantitative analysis of how countries perform in relation to a set of key socio-ecological predictors of biological invasions and of the capacity of countries to mitigate their impacts. Although economic and environmental factors are often considered important and are well-understood, we show that societal, technological and especially political factors are also essential for obtaining a comprehensive perspective on spatial and temporal changes in biological invasions. As expected from other studies (10, 14, 25–28), Trade was consistently the best predictor of EAS richness in one-predictor models, whereas the combination of Trade and Governance as main effects best explained EAS richness for most taxa in two-predictor models. These two predictors capture different aspects of biological invasions. Trade can facilitate the transportation of propagules and is therefore primarily linked to the introduction stage of biological invasions (29). In contrast, Governance is related to all invasion stages, from introduction to establishment and spread of alien species, as it is a proxy for the capacity and willingness to design and implement adequate policies to prevent alien species from transiting from one stage to the other. Nonetheless, Governance appears to limit biological invasions at high levels only (Figure 1). This likely reflects the complex interactions between Governance and other factors, including that awareness and willingness to respond decisively to biological invasions may increase only once substantial negative impacts of IAS have been widely observed in a country.

Lifestyle and Education is another predictor that proved to be important in our analyses but has been largely neglected so far. Lifestyle and Education was the best predictor of EAS richness for mammals and spiders when considering the 2015 data only. Lifestyle and Education was calculated by averaging the educational level of the population, the information globalization index and the cultural globalization index. Doing so enabled us to capture the potential level of understanding of complex issues such as biological invasions, but also connections with other cultures and countries, and the perception of nature (Table 1). Lifestyle and Education therefore has implications for alien species dispersal and establishment, e.g. via recreational activities and tourism, or mode of consumption. Importantly, Lifestyle and Education was also the best predictor for proactive national capacity and a good predictor for reactive national capacity. It is difficult at this stage to explain if this relationship is only correlative (countries investing in the education of their populations also tend to implement environmental policies) or if there is a causal relationship (populations with high levels of education may vote for governments more inclined to design and implement environmental laws). Our results nonetheless show that factors related to education and likely environmental awareness of a population, are important for predicting EAS richness and how countries will assess and react to the impacts caused by IAS.

### Effects of historical legacies on current levels of biological invasions

Our results underscore that invasion debt plays a crucial role in explaining current levels of biological invasions (13). We found that historical data, where available (i.e. for Trade, Governance), consistently better explained current numbers of EAS than did recent data. Due to a lack of predictor data prior to 1996, we were not able to analyze if – and for how long – historical legacies extend beyond this time. Time lags may also occur for other predictors, for which historical data were not available.

Biological invasions are the result of a range of processes that operate at different stages of the invasion process (30). For instance, while new alien species are introduced in response to changes in propagule pressure, introduced species become naturalized in response to human-induced changes in the recipient region and societal responses (e.g. IAS management, legislation) are adopted in response to observed or anticipated negative impacts (e.g. 24). These processes may be associated with substantial lag times: newly introduced species are often detected after a recording lag (31, 32), as does their spread to new locations and conversely, the adoption of effective management (33). Similarly, our findings show that historical levels of Governance, which are essential for the design and implementation of policies and the management of IAS, have an imprint on current EAS richness in countries. In particular, countries with higher levels of Governance 20 years ago tended to be less invaded than countries with intermediate Governance. Complex interactions between predictors suggest that historical legacies may also apply to other predictors. For example, since Lifestyle and Education was the most important predictor for explaining proactive capacities of countries to address issues related to IAS (Figure S4), its relationship with EAS richness is likely to be subject to time lag. Past Lifestyle and Education may also be a good predictor of current EAS richness, and it will likely be highly important for shaping future trajectories of EAS richness, as policies and management actions can take time to have effect.

### A global picture of country positions in the socio-economic space and implications for invasive alien species management and policies

Analyses of recent historical trajectories show that Trade has been increasing for all countries and will likely continue to do so in the next decades, with global freight demands predicted to increase three- to seven-fold between 2015 and 2050 (34, 35). Recent research has shown than under a business as usual scenario, we can expect a global increase in EAS richness of 36% between 2005 and 2050. The intensification of Trade will necessarily be followed by large increases in species introductions, and may therefore cause EAS richness increase to largely exceed the business as usual estimations.

For Governance, recent historical trajectories are much less uniform across regions and countries. In particular, there are strong differences between different regions of the world, with increases for some regions, such as non-EU Europe and Asia, and declines for others, such as Central America and Southern Africa (Figure 4). Differences are even larger at the country level and future country-specific projections for biological invasions, which are currently missing, would likely be highly uncertain. Overall, Governance appears to have an effect on EAS richness at high levels only. Among geopolitical regions whose level of Governance increased between 1996 and 2018 (Figure 4a), increases appear to be insufficient to reach the level of Governance at which it has an effect. Worse, the level of Governance of most geopolitical regions stagnated or even decreased over this period. Unless this trend is reversed, this will likely exacerbate the establishment of alien species whose rate of introduction will have also been increased by increases in Trade.

Our results nonetheless show that countries strongly differ regarding socio-ecological predictors of invasions. All the predictors we quantified in these analyses are related to different aspects of biological invasions and can therefore influence the future state of biological invasions (10, 17). Although understanding the interactions between these predictors is beyond the scope of this publication, this implies that there are substantial opportunities for countries to mitigate the impacts of biological invasions in the future (e.g. identifying predictors with the largest leverage or the potential to improve country ability to address biological invasions). Given the time lags involved in biological invasions, and the historical legacies of socio-ecological predictors on EAS richness, delays in positive changes, especially concerning Governance, may result in important long-term consequences for biodiversity.

Scenarios on biodiversity change to inform decision-making are under development (36, 37), but biological invasions are not considered in these analytical frameworks, despite the recognition of the importance of their integration into global environmental policies (e.g. Sustainable Development Goals, (38)). The on-going discussion on global targets for biodiversity conservation for the decades to come, including revised and specific targets on biological invasions (39), highlights that integrating biological invasions into thematically broad assessments of environmental change is crucial. By revealing that large increases in levels of Governance are required to mitigate increases in EAS richness resulting from the expected intensification of Trade, and identifying the regions of the world where such changes are critically needed, our socio-economic space for biological invasions paves the way for such integration.

## Methods

### Predictor selection and data

Based on previous findings (10, 17), we considered five main predictors (Table 1): i) Governance, i.e. the capacity of a country to design and implement policies, including policies aimed at addressing biological invasions; ii) Trade, as the most important predictor of propagule pressure; iii) Environmental Performance Index, i.e. the level of sustainable use of abiotic and biotic components of the recipient environment, including land use; iv) Lifestyle and Education, i.e. factors influencing people’s values and perception of nature, their understanding of the issue and their connections with other cultures and countries, with implications for alien species dispersal and establishment, e.g. via recreational activities and tourism, or mode of consumption; and v) Innovation, i.e. technological progress which can enhance the knowledge and technological means to manage biological invasions. To quantify each predictor, we searched for data available at the country scale from open access repositories with good transparency about the methods used to collate these variables, to ensure data quality and long-term maintenance. This resulted in a total of 12 variables extracted from the World Bank data repository (40), the KOF Swiss Economic Institute (41, 42), the Global Innovation Index (43) and the Wittgenstein Centre for Demography and Global Human Capital (44) (Table 1).

We extracted data on the selected variables for 2015, as this year corresponded to the final year for which data of the response variable, EAS richness, have been considered in our data set (see below). When data for this year were not available for a country, we used data from the most recent preceding year until 2010. To explore potential legacies of historical predictor conditions, we extracted historical data for Governance and Trade for each year from 1996 onwards, which was the first year for which these data were available for Governance; for the other predictors, data were available only for the more recent history. Altogether, predictor data were available for 125 countries (excluding some regions separate from mainland, which can have different invasion dynamics), which were then considered in the analyses (see Figure S1).

Further, following (15) we extracted mean annual temperature (BIO1) and mean annual precipitation (BIO12) for the years 1960 to 1990 from WorldClim (www.worldclim.org); for each country, we calculated mean annual temperature and total annual precipitation as the mean of raster cells within country borders. To control for area as well as sampling effect, we included country area (40) and sampling effort as additional predictor variables in our models. Sampling effort was measured using the metric proposed by (45), which is based on the number of GBIF records per unit area and accounting for native species richness. For reptiles, fishes and spiders, taxon-specific sampling effort was not available.

### Established alien species richness data

We calculated country-specific levels of invasion based on data of EAS richness of eight taxonomic groups for which global distribution data were available (plants, ants, amphibians, reptiles, fishes, birds, mammals and spiders) (15). Following (15), overall EAS richness was calculated by converting absolute EAS richness to a relative scale by dividing species richness by the maximum richness over all countries, resulting in values ranging from 0 to 1. Overall alien species richness for each country was then computed as the mean of relative richness values across taxonomic groups.

### National capacity data

Data representing countries’ capacity for reactive and proactive responses to IAS was obtained from (19). Proactive national capacity assesses the capacity of a country to prevent or early contain emerging incursions by IAS. Reactive national capacity accounts for the expertise, resources and willingness to mitigate the damage from IAS that are present in a country, which is essential to make IAS policy effective.

### Variable selection

The 12 socio-ecological variables selected to describe the main predictors of biological invasions (i.e. excluding climatic variables, country area and sampling effort) were interrelated in complex ways, resulting in collinearities. To keep predictors as independent from each other as possible and better disentangle their respective effects on the response variables described below, we imposed internal coherence between variables used to characterize a given predictor. To be coherent, variables characterizing a predictor had to be more correlated with each other than with variables characterizing other predictors. Variables that belong to a category but are more correlated with another likely indicate causal relationships between specific aspects of the two predictors that would cause a high correlation between predictors if they were included. Although understanding the causal relationships between predictors and the effects on biological invasions is interesting, this is beyond the scope of this study. Rather, maximizing independence between the predictors allows to better disentangle their respective effects on the response variables described below, whereas high correlations would lead to similar results in the analyses, rendering the analysis of the relationship difficult to interpret. We therefore discarded political globalization (initially considered to characterize Governance), which was more strongly correlated with imports (characterizing Trade) than with any of the other variables within its category (Figure S2); and per capita Gross National Income (initially considered to characterize Trade), which was more strongly correlated with control of corruption, government effectiveness and rule of law (characterizing Governance) than with imports. The remaining 10 socio-ecological variables were standardized to mean zero and unit standard deviation and then averaged per predictor for each country and year. In doing so, we avoided potential collinearity issues, reduced complexity and facilitated the interpretation of results (46).

### Analyzing the relationships between predictors and established alien species richness

We investigated the relationship between the five predictors and EAS richness per country with linear mixed-effects models (LMMs) using the lme function from the nlme R package v.3.1 (47, 48). To statistically identify non-linearities observed in preliminary analyses using splines, we fitted linear, second-order (quadratic) and third-order (cubic) models for each individual predictor. Quadratic models enabled us to detect accelerating (i.e. positive coefficients) or decelerating (i.e. negative coefficients) relationships. Similarly, we used cubic models to identify both accelerating and decelerating relationships across the range of values for a predictor.

We also incorporated mean annual temperature, total annual precipitation, mainland or island status of the country (represented by a categorical variable), as well as country area, sampling effort (ln-transformed) and their interaction (or only country area when sampling effort was not available for a taxonomic group) as fixed effects. We used overall EAS richness (ln-transformed to satisfy assumptions of normality of residuals and variance homogeneity) and EAS richness of each taxonomic group individually (ln [EAS richness + 1] transformed) as response variables. To account for spatial autocorrelation, we used broad geographical regions (level 2 nested in level 1 of the Biodiversity Information Standards – TDWG, (45)) as random effects. Alternative generalized linear mixed models using binomial and Poisson link functions on untransformed response variables provided qualitatively similar results (not shown), but could not be tested for spatial autocorrelation due to long computation times.

For each predictor X, we therefore assessed the following three models:

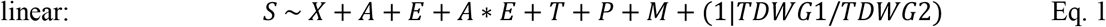

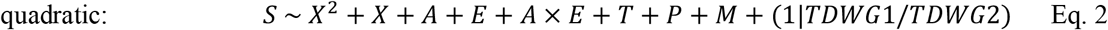

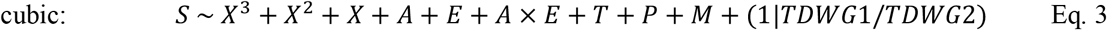

where S is EAS richness, X is a socio-economic predictor, A is country area, E is sampling effort (A was used instead of A + E + A × E for reptiles, fishes and spiders, for which sampling effort was not available), T is mean annual temperature, P is mean annual precipitation, M is mainland or island status and TDWG1 and TDWG2 are the levels 1 and 2 of the Biodiversity Information Standards. To avoid issues of data dredging, and because the focus of this study is on socio-economic predictors, polynomials were not tested on the spatial and climatic variables.

We assessed model performance using AICc (49), computed with the AICc function in the AICcmodavg R package v2.3-1 (50), and using the marginal variance explained after accounting for random effects, computed with the r^2^_nakagawa function in the performance R package v0.6.1 (51). We reported the model with the lowest AICc value for each predictor (due to the large number and variety of models compared, we do not report all ΔAICc values).

We also computed LMMs incorporating both Governance (G) and Trade (Tr) in the models (using the linear, quadratic and cubic transformations for both predictors; Eqs 4–6).

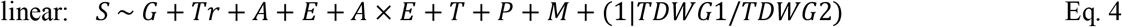

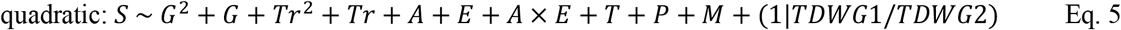

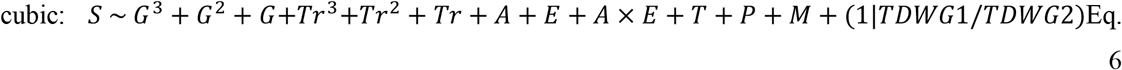

Finally, for models using Governance and Trade as predictors, we performed analyses for historical predictor conditions for 1996 and for the annual values averaged between 1996 and 2015 (historical data were not available for the other three predictors). The same 125 countries were used in all analyses, permitting comparison with respective models using the 2015 data. A lower AICc value for models using 1996 data than using 2015 data would reveal the historical legacy of these predictors on EAS richness. Alternative models

In these analyses, we did not combine other predictors than Governance and Trade due to high collinearity (see results for values). Incorporating additional predictors in exploratory analyses led to Variance Inflation Factors > 3 in the models (results not shown). Note also that we did not use the axis of the PCA described above as predictors for two reasons: i) that would have prevented the exploration of the effects of historical data due to lack of data for other predictors; ii) that enabled us to better explore the effects of the different predictors on the different response variables we considered (see national response capacities below).

### Analyzing the relationships between predictors and national response capacities

We analyzed the relevance of the five main predictors of invasions for the ability of countries to control and manage biological invasions, i.e. their national response capacities (19). We modified Eqs 1–6 using proactive and reactive national capacities as response variables and removed the biological (T, P, M) and statistical (A, E) predictors of EAS richness (Eqs 7–12). We hypothesized that Governance, Environmental Performance and Lifestyle and Education should be positively correlated with proactive capacity of a country to prevent or rapidly respond to emerging incursions by IAS. As Trade is expected to lead to more species introductions (14), which in turn should lead to more reactive measures due to rising awareness of the impacts of IAS, we argued that Trade should show a stronger correlation with the reactive capacity of countries to mitigate negative impacts caused by IAS already present. As for EAS richness, models were evaluated with current (2015) and historical (1996) predictor data, and averaged over the 1996–2015 period.

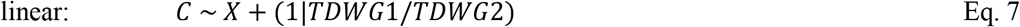

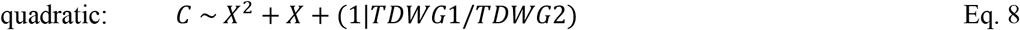

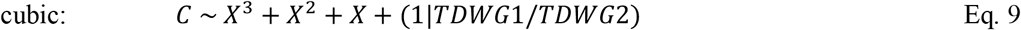

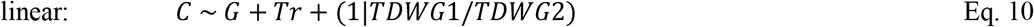

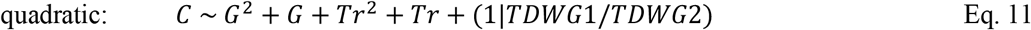

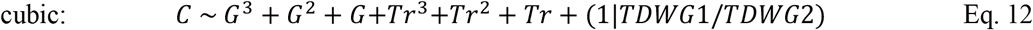

where C is proactive or reactive national capacity and the other notations are as in Eqs 7–12.

For all models, we tested for residual spatial autocorrelation by constructing correlograms of Moran’s I in relation to increasing distance between country centroids using the spline.correlog function in the ncf R package v1.2-9 (52). Significance was assessed using 95% confidence intervals, built from 1000 bootstrapped randomizations of the residuals (Figures S6, S7). All statistical analyses were performed with the R software v. 4.0.3 (48).

### Visualization of countries in a two-dimensional socio-economic space

We mapped countries in a two-dimensional space defined by the 2015 levels of Governance and Trade. To facilitate the interpretation of results, countries were assigned to different geopolitical regions. To identify groups of countries that differ distinctly from each other, we applied two hierarchical clustering algorithms based on distance between countries in this socio-economic space. We used the complete-linkage and the Ward methods in the R function hclust from the default stats package. To evaluate the number of clusters best separating the countries, we used the function NbClust from the NbClust R Package v3.0 (53), which evaluates the number of clusters based on 30 different indices.

We used data from 2015 in the previous analyses because this corresponds to the most recent year for which EAS richness and national response capacity data were both available; as data for Governance and Trade were available until 2018, we mapped countries for the full range of data when investigating their trajectories in the socio-economic space through time.

## Supporting information

Data for mixed models

Data for temporal trends

## Data Availability

All data analyzed here are freely available from the original sources provided in Table 1. The data used as predictors for the three time periods (1996, 1996-2015, 2015) have been compiled in a single CSV file available in supplementary material.

## Acknowledgements

This work was funded by the BiodivERsA-Belmont Forum Project “Alien Scenarios” (GL, BL, SD, DM, FE: FWF project no I 4011-B32; NRP, CPG, EC: Project PCI2018-092966, funded by FEDER/Ministerio de Ciencia e Innovación – Agencia Estatal de Investigación; MG and IK: BMBF/PT DLR 01LC1807C; HS: BMBF/PT DLR 01LC1807A; JMJ: BMBF/PT DLR 01LC1807B; FC, CB, CD, AT: ANR-18-EBI4-0004).

