## Supplementary Information for "Capacity of countries to reduce biological invasions"

**Supplementary Material**

Guillaume Latombe^ab,1^, Hanno Seebens^c^, Bernd Lenzner^a^, Franck Courchamp^d^, Stefan Dullinger^a^, Marina Golivets^e^, Ingolf Kühn^efg^, Brian Leung^h^, Núria Roura-Pascual^i^, Emma Cebrian^jk^, Wayne Dawson^l^, Christophe Diagne^d^, Jonathan M. Jeschke^mno^, Cristian Perez-Granados^ip^, Chunlong Liu^dmno^, Dietmar Moser^a^, Anna Turbelin^d^, Piero Visconti^q^, Franz Essl^a^

^a^BioInvasions, Global Change, Macroecology-Group, Department of Botany and Biodiversity Research, University Vienna, Rennweg 14, 1030 Vienna

^b^Institute of Evolutionary Biology, The University of Edinburgh, King’s Buildings, Edinburgh EH9 3FL, United Kingdom

^c^Senckenberg Biodiversity and Climate Research Centre, Senckenberganlage 25, 60325 Frankfurt, Germany

^d^Université Paris-Saclay, CNRS, AgroParisTech, Ecologie Systématique Evolution, 91405 Orsay, France

^e^Helmholtz Centre for Environmental Research – UFZ, Theodor-Lieser-Str. 4, 06120 Halle, Germany

^f^Martin Luther University Halle-Wittenberg, Geobotany and Botanical Garden, Halle, Germany

^g^German Centre for Integrative Biodiversity Research (iDiv) Halle-Jena-Leipzig Deutscher Platz 5e, 04103 Leipzig, Germany; ORCID: 0000-0003-1691-8249

^h^Department of Biology. McGill University. Montreal, Quebec, Canada. H3A 1B1.

^i^Departament de Ciències Ambientals, Facultat de Ciències, Universitat de Girona, Girona, Catalonia.

^j^Centre d’Estudis Avançats de Blanes-CSIC, Girona, Spain;

^k^GRMAR, Institute of Aquatic Ecology, University of Girona, Girona, Spain

^l^Department of Biosciences, Durham University, South Road, Durham, DH1 3LE, UK

^m^Institute of Biology, Freie Universität Berlin, 14195 Berlin, Germany

^n^Leibniz Institute of Freshwater Ecology and Inland Fisheries (IGB), 12587 Berlin, Germany

^o^Berlin-Brandenburg Institute of Advanced Biodiversity Research (BBIB), 14195 Berlin, Germany

^p^Ecology Department. Universidad de Alicante, 03080. Alicante. Spain

^q^Biodiversity, Ecology and Conservation Group, International Institute for Applied System Analyses. A-2361 Laxenburg, Austria

**Table S1.** Results of model selection for explaining EAS richness and national capacities in 125 countries based on the small-sample size corrected Akaike Information Criterion (AICc). Shown are the models with the lowest AICc values. The ΔAICc is the difference with the lowest 2015 values. r^2^ values are the marginal variance. An asterisk (*) indicates results compared to Governance and Trade values only for 2015. Values between brackets indicate the gains in marginal r^2^ compared to models with no predictor.

| **Response variable** | **Predictors** | **Model type** | **Time period** | **ΔAICc** | **Marginal r^2^** |
| --- | --- | --- | --- | --- | --- |
| All taxa combined | Trade | Quadratic | 1996 | -19.02 | 0.83  (+0.04) |
| Plants | Trade | Quadratic | 1996-2015 | -2.13 | 0.56  (+0.02) |
| Ants | Trade + Governance | Quadratic | 1996-2015 | -2.11 | 0.62  (+0.01) |
| Amphibians | Trade | Quadratic | 1996 | -9.81 | 0.65  (+0.06) |
| Reptiles | Trade + Governance | Cubic | 1996-2015 | -6.70 | 0.51  (+0.05) |
| Fishes | Trade + Governance | Quadratic | 1996 | -6.68 | 0.60  (+0.04) |
| Birds | Trade + Governance | Quadratic | 1996 | -21.14 | 0.76  (+0.06) |
| Mammals | Trade + Governance | Quadratic | 1996 | -5.12 | 0.67  (+0.07) |
| Spiders | Trade + Governance | Cubic | 1996 | -5.14 | 0.64  (+0.05) |
| Proactive national capacity | Lifestyle and Education | Quadratic | 2015 | 0.00 | 0.47 |
| Proactive national capacity* | Governance | Quadratic | 1996-2015 | 4.91  (-1.01) | 0.33  (+0.02) |
| Reactive national capacity | Trade | Quadratic | 1996 | -2.16 | 0.28  (+0.02) |

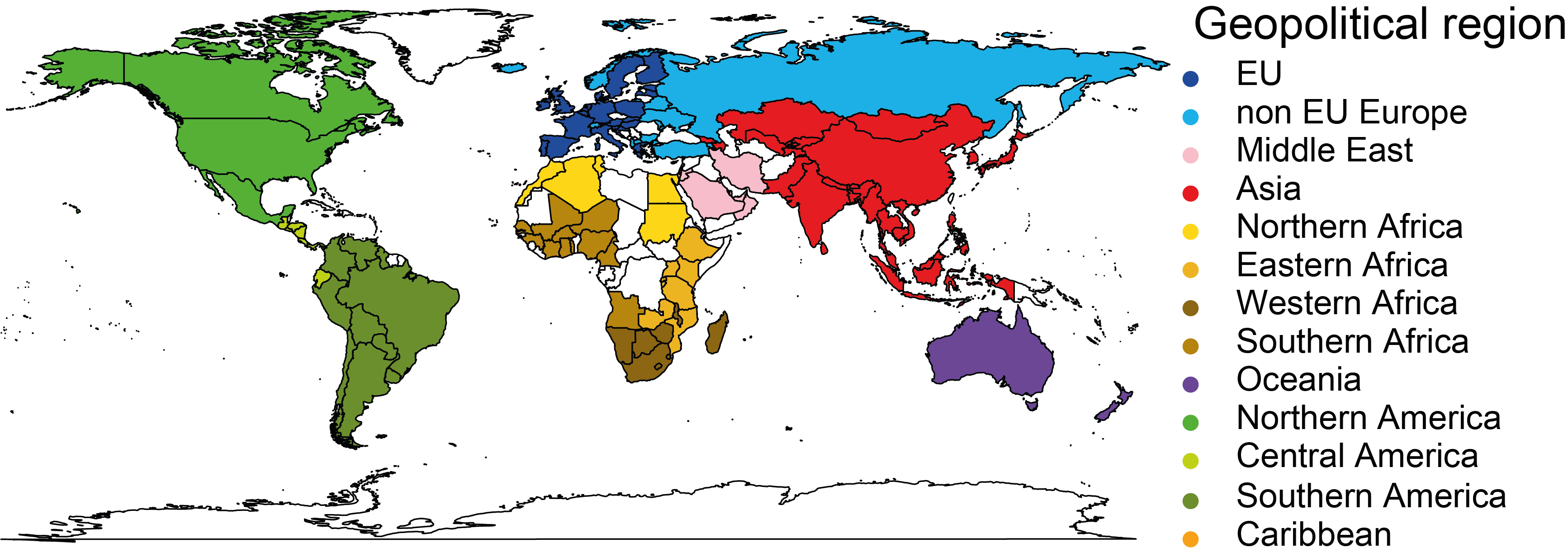

**Figure S1.** Geopolitical regions of the 125 countries (excluding some regions separate from mainland) included in the analyses. Countries in white were not considered due to data deficiency.

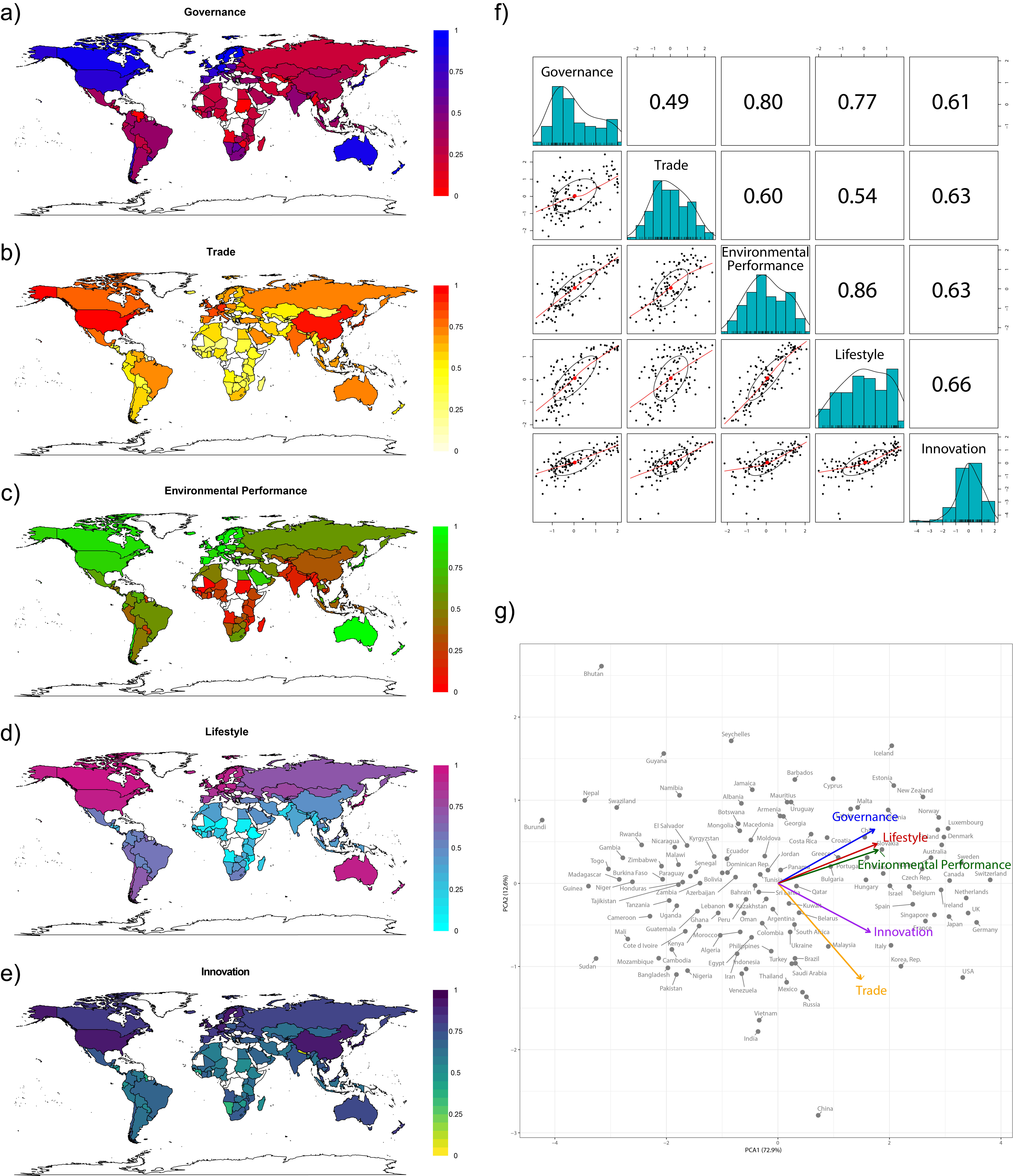

**Figure S2.** Relationships between five predictors and the respective variables (i.e. economy: Trade, policy: Governance, environment: Environmental Performance, social norms: Lifestyle and Education, technology: Innovation) of biological invasions used in the analyses. a-e) The 125 countries (excluding some regions separate from mainland) used in the analyses are shown in different colors based on their levels for each of the five predictors. f) Pairwise Pearson's r correlation analyses between the predictors. h) Principal component analyses of the five predictors (blue: Governance, orange: Trade, green: Environmental Performance, red: Lifestyle and Education, purple: Innovation).

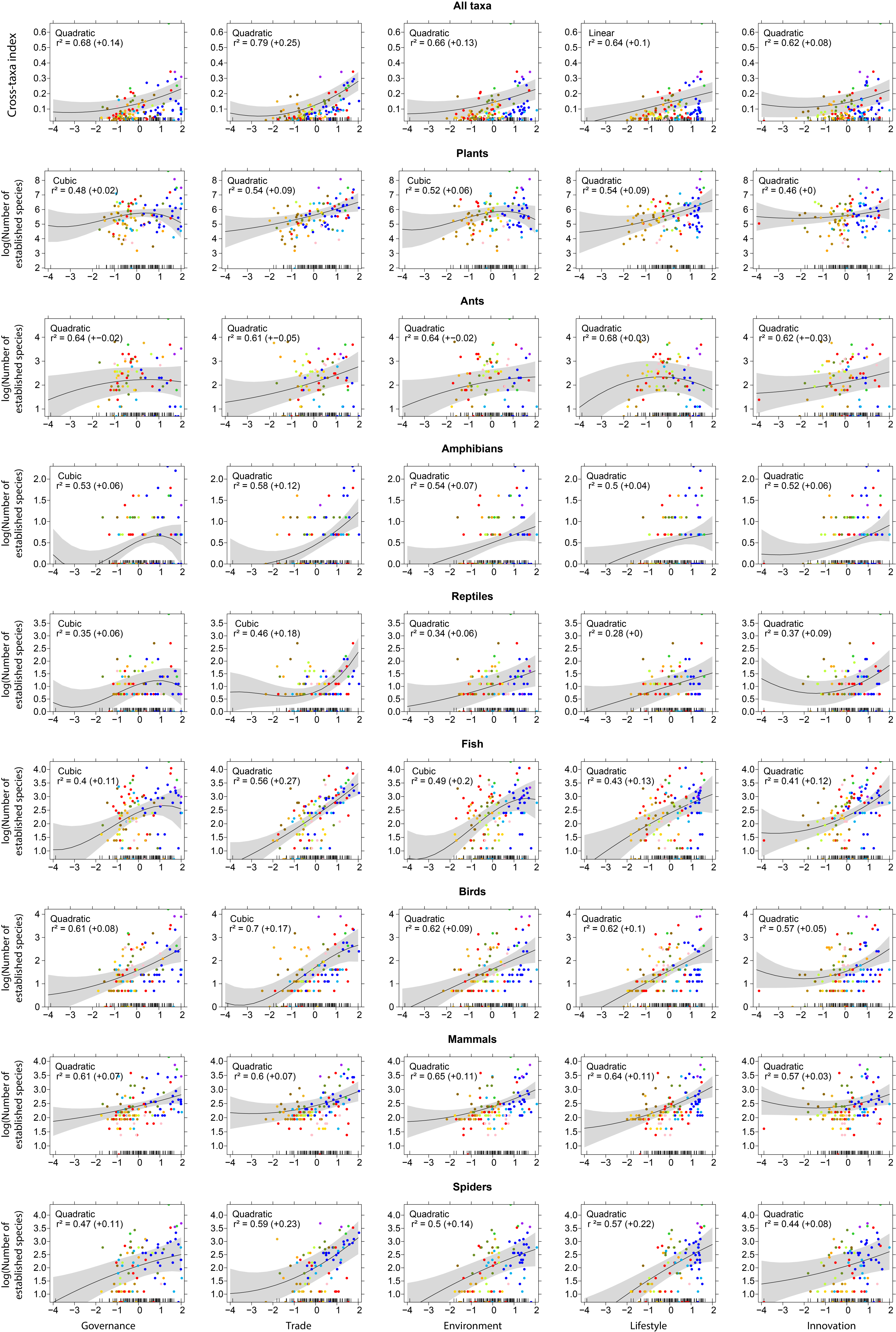

**Figure S3.** Relationships between the five predictors and overall EAS richness. The number of EAS was controlled for by country area, sampling effort, mean annual temperature and mean annual precipitation. The type of regression displayed is the one with the lowest AICc. The colors represent the geopolitical regions that the countries belong to (for legend, see Figures 1 and S1).

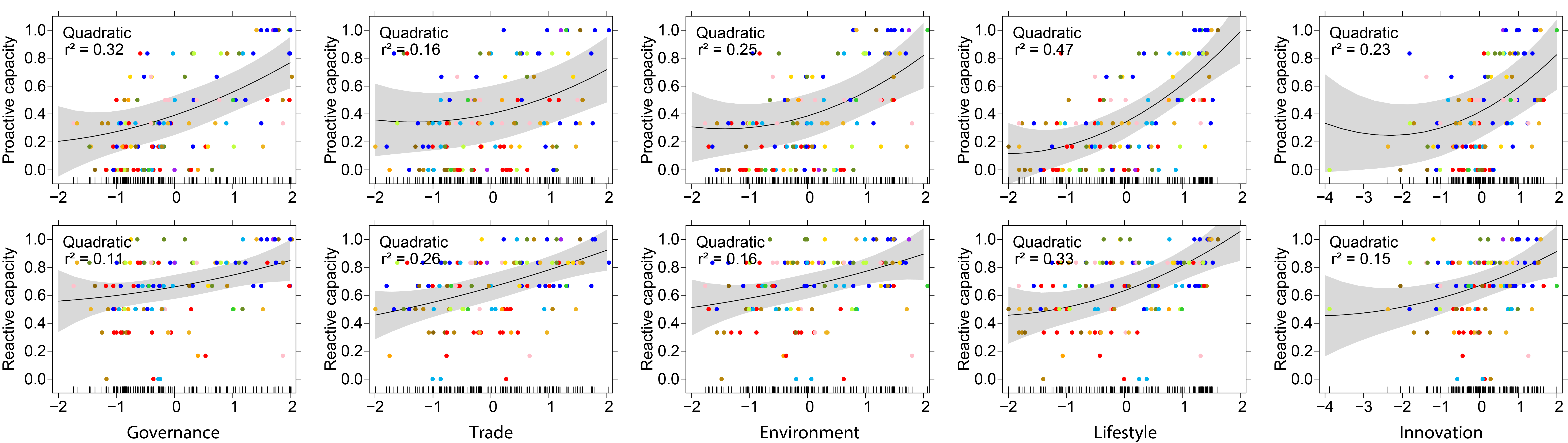

**Figure S4.** Relationships between the five predictors (2015) and national capacities. The type of regression displayed is the one with the lowest AICc. The colors represent the regions the geopolitical countries belong to (for legend, see Figures 1 and S1).

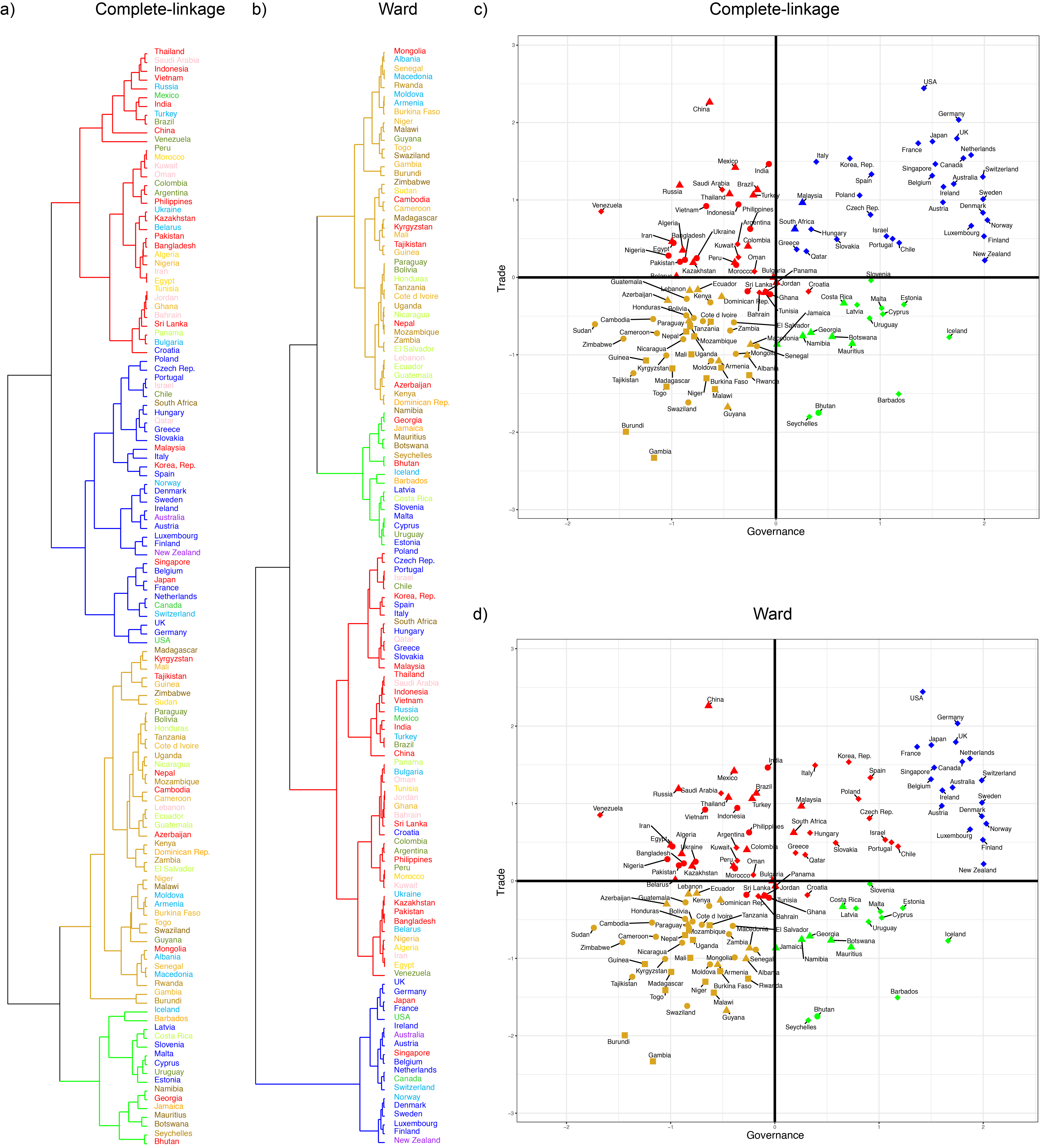

**Figure S5.** Cluster analysis of 125 countries in the socio-economic space based on Governance and Trade. a,b) Dendrograms of 125 countries based on their position in the two-dimensional socio-economic space using two different cluster algorithms (i.e. complete-link and Ward). The colors of the country names represent the geopolitical regions (see Fig. S1). The colors of the branches represent the cluster countries belong to. d,e) Countries in the socio-economic space are colored according to the cluster they belong to.

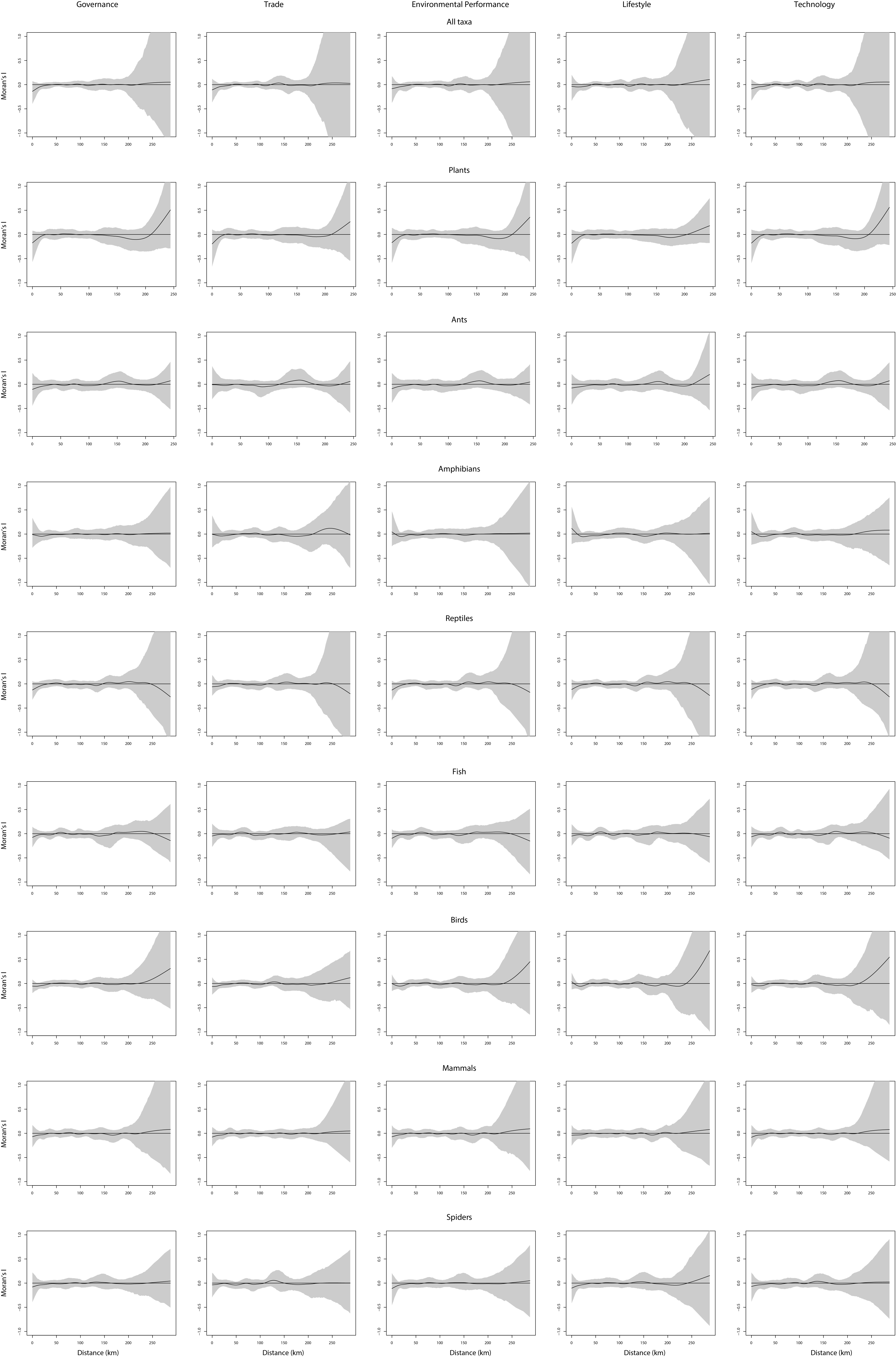

**Figure S6.** Moran’s *I* correlograms of residuals in the quadratic models for EAS richness. The X-axes show distance between country centroids. Spatial autocorrelation was always low at large and medium distances, and increases somewhat for some taxonomic groups at short distances, albeit not significantly.

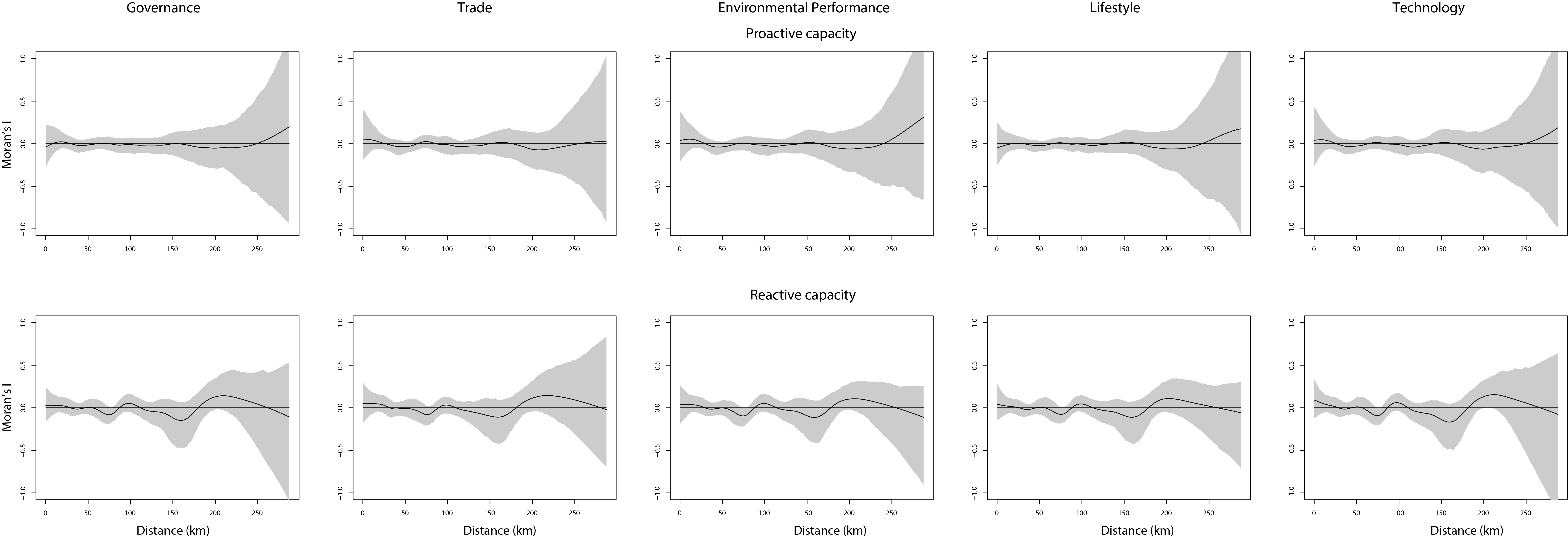

**Figure S7.** Analysis of spatial autocorrelation. Shown are correlograms of Moran’s *I* in relation to distance between country centroids for the quadratic models using recent predictor data (2015) to explain national capacities to mitigate negative impacts of biological invasions. Autocorrelation is low in all models and at all distances.
